# protein-sol pKa: prediction of electrostatic frustration, with application to coronaviruses

**DOI:** 10.1101/2020.04.21.053967

**Authors:** Max Hebditch, Jim Warwicker

## Abstract

Evolution couples differences in ambient pH to biological function through protonatable groups, in particular those that switch from buried to exposed and alter protonation state in doing so. We present a tool focusing on structure-based discovery and display of these groups. Since prediction of buried group pKas is computationally intensive, solvent accessibility of ionisable groups is displayed, from which the user can iteratively select pKa calculation centers. Results are color-coded, with emphasis on buried groups. Utility is demonstrated with coronaviruses, which exhibit variable dependence on the acidic pH of the endocytotic pathway. After benchmarking with variants of murine hepatitis virus, a pair of conserved histidine residues are identified that are predicted to be electrostatically frustrated at acidic pH in a common structural core of pre- and post-fusion coronavirus spike proteins. We suggest that an intermediate expanded conformation at endosomal pH could relax the frustration, allowing histidine protonation, and facilitating conformational conversion. This tool is available at http://www.protein-sol.manchester.ac.uk/pka/.

## Introduction

Since pKas underlie pH-dependent phenomena in biology, their prediction has received significant attention, largely through continuum electrostatics methods (Alexov *et al*., 2011). We have contributed a server for predicting pH and ionic strength dependence with a Debye-Hückel (DH) model that accounts for solvent exposed groups, which are generally in the great majority (Hebditch and Warwicker, 2019). However, conformational change often depends on the electrostatic frustration (destabilization) that develops when a buried group cannot ionize at a pH where it would in a more solvent accessible conformation (Narayan and Naganathan, 2018). We reasoned that a web tool focusing on buried ionizable groups would be useful for studying pH-dependent conformational change, and have adapted our existing mixed finite difference Poisson-Boltzmann (FDPB) and DH model. Here, the server is demonstrated with coronaviruses, some of which use the endocytotic pathway for membrane fusion, whereas others fuse at the plasma membrane. We focus on the pre- to post-fusion conformational changes in the S2 part of the spike protein, that mediates membrane fusion (Heald-Sargent and Gallagher, 2012).

## Methods

We have sought to limit FDPB/DH run time for pKa predictions (Warwicker, 2004) to about two minutes processing. Upon upload of a structure, the user is presented with a color-coded display (NGL viewer, Rose *et al*., 2018) of solvent accessible surface area (ASA) values for ionizable groups. A user iteratively specifies centers, around which pKa calculations are made for spheres of radius 25 Å (about the size of lysozyme). Edge effects do not have a big effect on predicted pKas towards the sphere centre. Results for Asp, Glu, Lys, Arg, His accumulate as more centers are added, and are color-coded to show whether a group is stabilizing or destabilizing, assessed from the difference between calculated and intrinsic pKa (capped at −5 and 5). Users may either use ionisable group ASA/burial or literature knowledge of interesting sites, to select pKa calculation centers. It is envisaged that the server will allow a user to quickly survey a set of structures for potential pH-dependence hotspots, rather than provide a great depth of analysis for each structure.

## Results

Although there are about 40 structures of pre-fusion coronavirus spike proteins, there is just one post-fusion structure available (April 2020) that extends beyond the helical fusion core, for murine hepatitis virus (MHV) strain A59 (6b3o, Walls *et al*., 2017). A pre-fusion structure for MHV A59 is also available (3jcl, Walls *et al*., 2016). In a variant of mouse hepatitis virus type 4 (MHV4), spike protein Q1015H, Q1042H (MHV A59 numbering) and one further mutation (L to R) render the virus pH-dependent, via the endocytotic pathway (Gallagher *et al*., 1991). Modelling the mutations to histidine in the MHV A59 background, we find they are predicted to be buried in the pre-fusion form (destabilizing, not shown) and exposed in the post-fusion structure (not destabilizing, Fig. 1a). Color schemes for ASA and pKa-related stabilization are given (Fig. 1b). Our results are consistent with the relief of electrostatic frustration at endosomal pH biasing conformation away from the pre-fusion structure. The story is a little more complicated, since the MHV4 variant also loses fusion activity at neutral pH (Gallagher *et al*., 1991), which could be due to additional stabilization of the pre-fusion form with pH-independent histidine interactions.

**Fig. 1.**
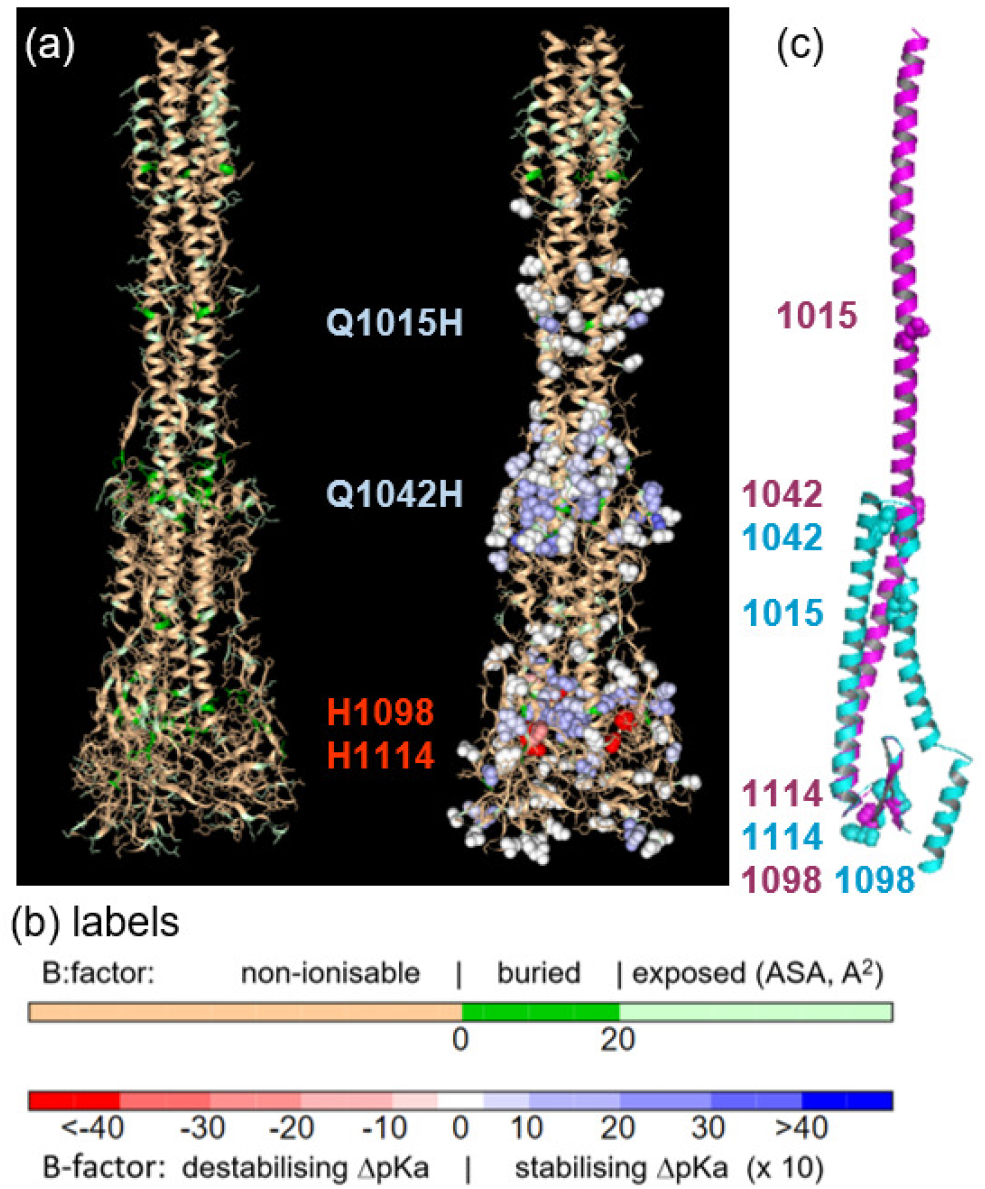
The pKa web server. (a) The post-fusion structure of MHV A59 (6b3o) with ionizable group burial (left) and pKa calculations around selected centres (right), following color codes in (b). (c) Segments of pre- (3jcl, cyan) and post-fusion (6b3o, magenta) MHV A59 structures. Residues of interest are indicated throughout.

Fig. 1c shows equivalent parts (972 – 1118) of a monomer from pre- and post-fusion MHV A59, structurally aligned through a small common core around 1098/1114. Extending from the structurally aligned segments are helices that demonstrate the extensive spike protein structural changes that go along with cell fusion. Whereas the helical region wraps back around the aligned core in the pre-fusion structure, it extends upwards post-fusion, carrying the fusion peptide towards its target membrane, shown by the relative locations of 1015. Switching to conserved histidines and general coronavirus features, only two are present across the spike proteins of coronaviruses, H1098 and H1114 (MHV A59 numbering). In 37 of 38 pre-fusion coronavirus spike protein structures, as well as the post-fusion structure, H1098 and H1114 are buried and predicted to be destabilizing upon exposure to acidic pH (Fig. 1a, red spacefill). If these conserved histidines are electrostatically frustrated in both pre- and post-fusion conformations at endosomal pH, they would not bias towards either form. However, to allow the extensive changes exemplified in Fig. 1c, it is possible that the core region around H1098/H1114 loosens. If H1098 and/or H1114 were solvent exposed and protonated, then relief from frustration in a conformational intermediate could play a role in facilitating transfer between post- and pre-fusion structures in coronaviruses that use the endocytotic pathway, including SARS-CoV-2 (Ou *et al*., 2020). In this proposal, H1098/H1114 assistance in crossing the pre- to post-fusion conformational barrier would be available to those viruses that are unable to fuse at the plasma membrane. Interestingly, both H1098A and H1114A mutations in MHV A59 prevented virus growth (Li *et al*., 2018), perhaps indicative of (pH-independent) packing stabilizations in their buried environments, so that evolutionary retention for fusion could be afforded by a more direct structural imperative. This would be in line with the coupling of factors that determine infection pathways, including spike protein stability, receptor binding, proteolytic cleavage, as well as endosomal pH (Heald-Sargent and Gallagher, 2012; Millet and Whittaker, 2015). If the proposed loosening around the conserved post- and pre-fusion core is borne out, then although it may not be universally necessary (in pH-dependent entry), it could be the basis for a novel, albeit transient, coronavirus drug target. Our web tool will allow users to look for ionisable groups that could mediate pH-dependence in coronaviruses and other systems.

## References

Alexov, E. et al. (2011) Progress in the prediction of pKa values in proteins. Proteins, 79, 3374–3380.

Gallagher, T.M. et al. (1991) Alteration of the pH-dependence of coronavirus-induced cell fusion: effect of mutations in the spike glycoprotein. J. Virol., 65, 1916–1928.

Heald-Sargent, T. and Gallagher, T. (2012) Ready, set, fuse! The coronavirus spike protein and acquisition of fusion competence. Viruses, 4, 557–580.

Hebditch, M. and Warwicker, J. (2019) Web-based display of protein surface and pH-dependent properties for assessing the developability of biotherapeutics. Sci. Rep., 9, 1979.

Li, P. et al. (2018) Identification of H209 as essential for pH8-triggered receptor-independent syncytium formation by S protein of mouse hepatitis virus A59. J. Virol., 92, e00209–18.

Millet, J.K. and Whittaker, G.R. (2015) Host cell proteases: critical determinants of coronavirus tropism and pathogenesis. Virus Res., 202, 120–134.

Narayan, A. and Naganathan, A.N. (2018) Switching protein conformational states by protonation and mutation. J. Phys. Chem. B., 122, 11039–11047.

Ou, X. et al. (2020) Characterization of spike glycoprotein of SARS-Cov-2 on virus entry and its immune cross-reactivity with SARS-CoV. Nat. Comm., 11, 1620.

Rose, A.S. et al. (2018) NGL viewer: web-based molecular graphics for large complexes. Bioinformatics, 34, 3755–3758.

Walls, A.S. et al. (2016) Cryo-electron microscopy structure of a coronavirus spike glycoprotein structure. Nature, 53, 47–52.

Walls, A.S. et al. (2017) Tectonic conformational changes of a coronavirus spike glycoprotein promote membrane fusion. Proc. Natl. Acad. Sci. USA, 114, 11157–11162.

Warwicker, J. (2004) Improved pKa calculations through flexibility based sampling of a water-dominated interaction scheme). Protein Sci., 13, 2793–2805.

